## Supplementary Materials for "Multiple pathways for glucose phosphate transport and utilization support growth of *Cryptosporidium parvum*"

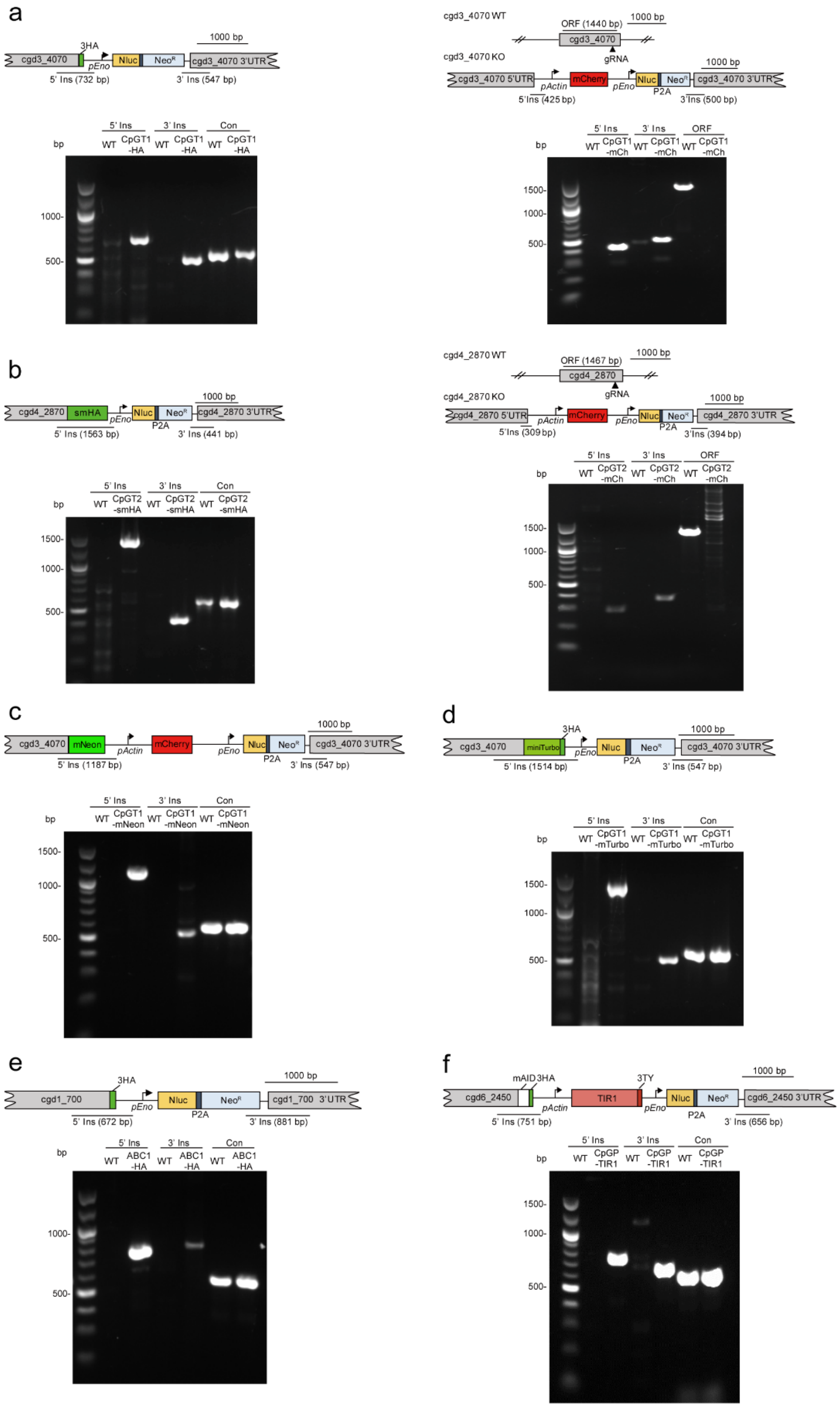


**Supplementary Fig. 1: Maps of the *C. parvum* genes modified in this study and confirmation by PCR analysis**

**a** Maps of native and modified loci of CpGT1. PCR analysis demonstrated successful insertion into the CpGT1 locus for epitope tagging with HA (left) and complete deletion (right) with predicted 5’ and 3’ insertion sites. **b** Maps of native and modified loci of CpGT2. PCR analysis demonstrated successful insertion into the CpGT2 locus for epitope tagging with smHA (left) and complete deletion (right) with predicted 5’ and 3’ insertion sites. **c** Maps of modified loci of CpGT1 containing mNeon as C-terminal fusion and cytosolic mCherry reporter driven by the *Cp* actin promoter. PCR analysis demonstrated successful insertion into CpGT1 locus with predicted 5’ and 3’ insertion sites. **d** Maps of modified loci of CpGT1 with C-terminal miniTurbo fusion. PCR analysis demonstrated successful insertion into CpGT1 locus with predicted 5’ and 3’ insertion sites with WT as the control. **e** Maps of modified loci of CpABC1 genes and PCR analysis demonstrated successful epitope tagging of the CpAB1 locus with predicted 5’ and 3’ insertion sites. **f** Maps of modified loci of CpGP gene with mAID-3HA at C-terminus and TIR-3TY driven by the *Cp* actin promoter. PCR analysis demonstrated successful insertion into CpGP locus with predicted 5’ and 3’ insertion sites. WT, wild type. Con, detects the 588-bp sequence from an unrelated gene (cgd5_4450). ORF, detects the open reading fragment of induvial gene. All PCR experiments were performed twice with similar results.

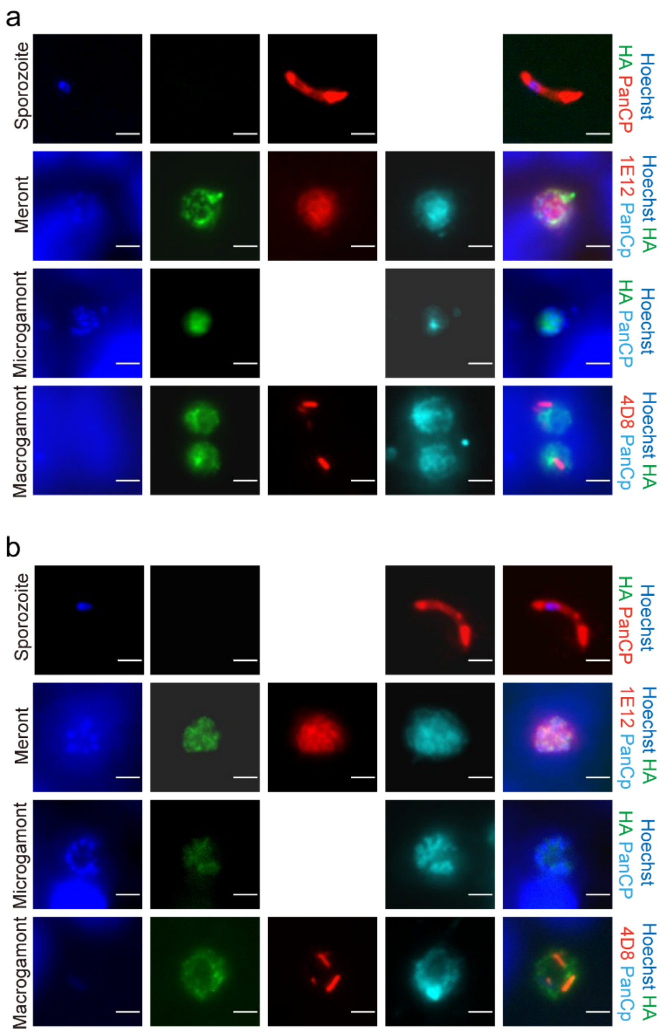


**Supplementary Fig. 2: Immunofluorescence localization of CpGT1-3HA (a) and CpGT2-smHA (b) in different life stages of *C. parvum***

To examine the pattern in extracellular parasites, CpGT1-3HA or CpGT2-smHA sporozoites were fixed, and stained with rat anti-HA (green), PanCp (red), Hoechst (blue). To examine the pattern in intracellular stages, HCT-8 cells were infected with CpGT1-3HA or CpGT2-smHA oocysts and fixed at 4 h (trophozoite), 24 h (meront), or 48 h (microgamont and macrogamont). For meront, coverslips were stained with rat anti-HA (green), mouse 1E12 (red) that localizes to the parasite membrane, rabbit PanCp (cyan), Hoechst (blue). For microgamont and macrogamont, coverslips were stained with rat anti-HA (green), mouse 4D8 (red) that localizes to macrogamont, rabbit PanCp (cyan), Hoechst (blue). The experiment was performed twice with similar results. Scale bars = 2 µm.

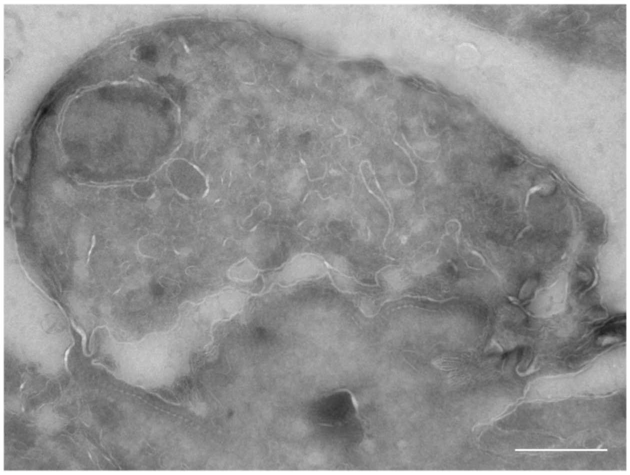

**Supplementary Fig. 3: Immuno-EM of WT *C. parvum***

Wild type *Cp* were grown in HCT-8 cells, fixed at 20 hpi and processed for immuno-EM and stained with rabbit anti-HA followed by 18-nm colloidal gold goat anti-rabbit IgG. The experiment was performed twice and cells were consistently negative for anti-HA staining. Scale bar = 500 nM.

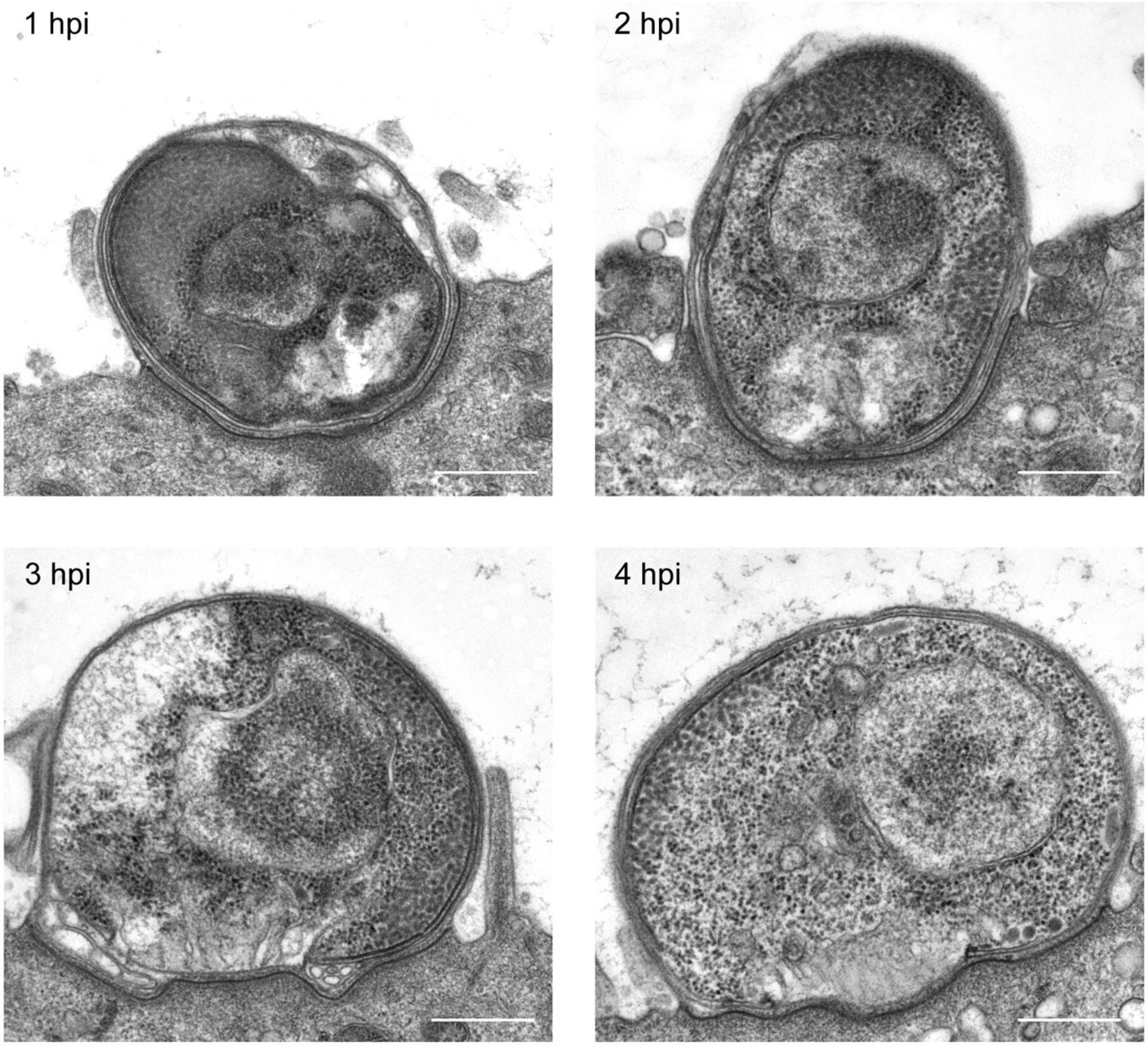


**Supplementary Fig 4. Transmission electron micrographs of feeder organelle formation followed by invasion**

Human intestinal spheroids were cultured on transwells to create the air-liquid interface culture using a modification of previously published methods [1,2]. Cells were infected with WT parasites and monolayers were fixed and processed at intervals post infection. Scale bars = 500 nm.

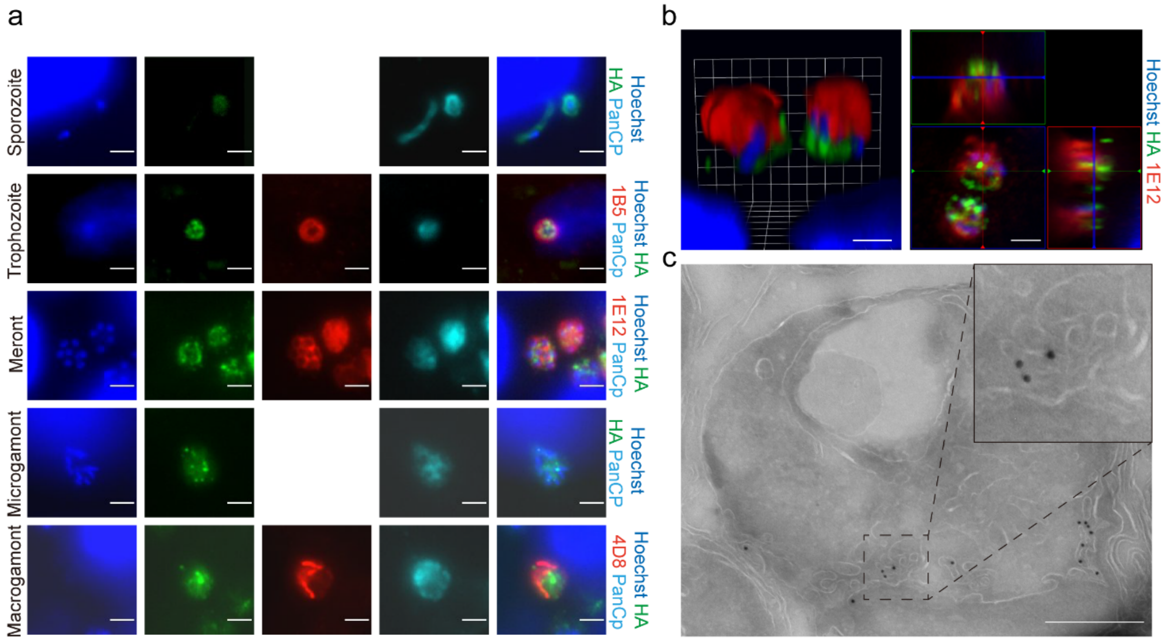


**Supplementary Fig. 5: CpABC1 was localized to the *C. parvum* feeder organelle.**

**a** Immunofluorescence of CpABC1-3HA in different life stages of *C. parvum*. HCT-8 cells were infected with CpABC1-3HA oocysts fixed at 4 h (sporozoites, trophozoite), 24 h (meront), or 48 h (microgamont and macrogamont). For sporozoites and trophozoite staining, monolayers were stained with rat anti-HA (green), mouse 1B5 (red) that localizes to the parasite-host interface, rabbit PanCp (cyan), Hoechst (blue). For meront staining, coverslips were stained with rat anti-HA (green), mouse 1E12 (red) that localizes to the parasite membrane, rabbit PanCp (cyan), Hoechst (blue). For microgamont and macrogamont staining, coverslips were stained with rat anti-HA (green), mouse 4D8 (red) that localizes to macrogamont, rabbit PanCp (cyan), Hoechst (blue). The experiment was performed twice with similar results. Scale bars = 2 µm. **b** Immunofluorescence of CpABC1-3HA in meront stage by confocal microscopy to generate a Z-stack that was rendered in 3 dimensions using Volocity software. HCT-8 cells were infected with CpABC1-3HA oocysts. After 24 h, monolayers were fixed and stained with rat anti-HA (green), mouse 1E12 (red), Hoechst (blue). The experiment was performed twice with similar results. Scale bars = 2 µm. **c** Transmission electron micrographs of the meront expressing CpABC1-3HA. HCT-8 cells were infected with CpABC1-3HA parasites. After 20 hpi, monolayers were fixed and stained with rabbit anti-HA followed by 18-nm colloidal gold goat anti-rabbit IgG. Scale bars = 500 nM.

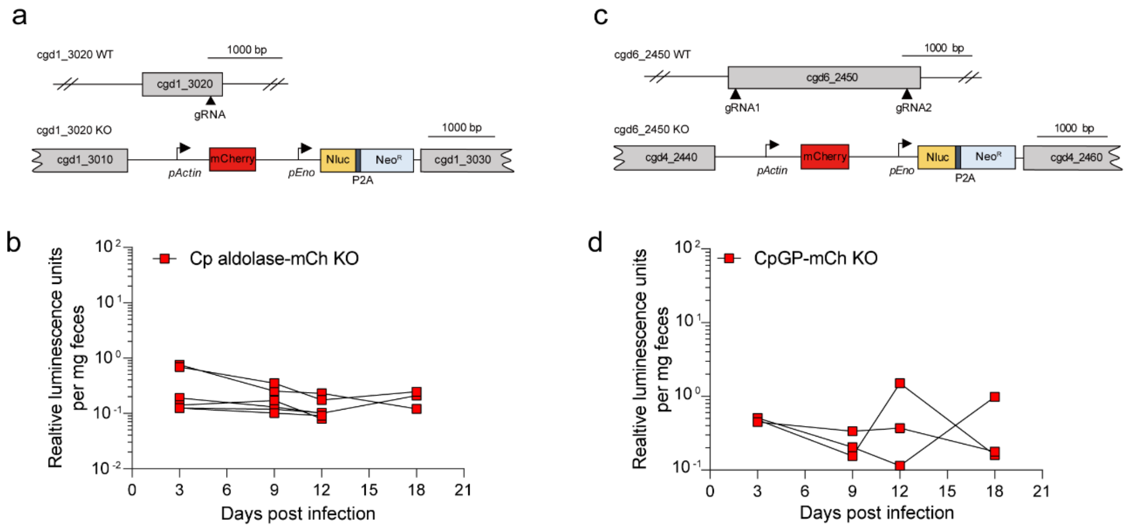


**Supplementary Fig. 6: Attempts to knockout *Cp* aldolase and *Cp* glucan phosphorylase (CpGP) in *C. parvum* were unsuccessful**

**a** Maps of native and modified loci of the *Cp* aldolase gene showing the strategy for generating a gene knockout. **b** Relative luminescence units per milligram feces. Nano-luciferase assays were performed on fecal pellets collected from GKO mice infected with transfected sporozoites. Each line represents an individual mouse (n = 6, from two independent experiments). **c** Maps of native and modified loci of the CpGP gene showing the strategy for generating a gene knockout. **d** Relative luminescence units per milligram feces. Nano-luciferase assays were performed on fecal pellets collected from GKO mice infected with transfected sporozoites. Each line represents an individual mouse (n = 3, from a single experiment).

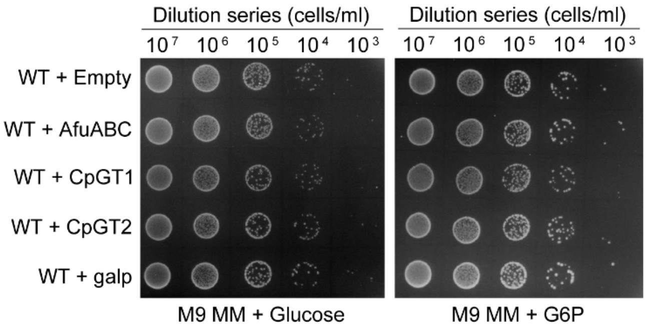


**Supplementary Fig. 7: Growth of wild type *E. coli* transformed expressing different transporters on glucose or glucose-6-phisphate (G6P)**

Growth of WT K-12 *E. coli* (BW25113) on M9 minimal media (M9 MM) agar plates supplemented with 10 mM glucose or glucose-6-phosphate (G6P). The indicated dilutions were plated and grown at 37 °C for 40 h before imaging. This experiment was performed three times with similar results.

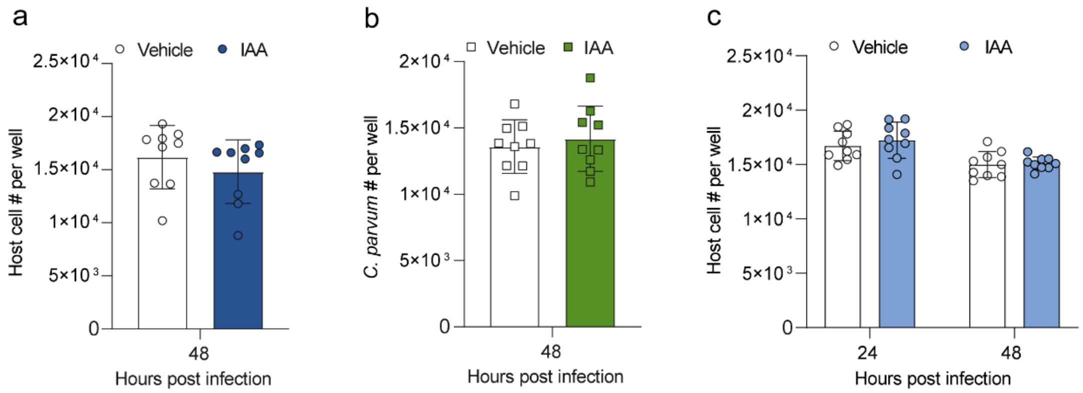


**Supplementary Fig. 8: Growth of *C. parvum* on HCT-8 cells in presence or absence of IAA**

**a, b** HCT-8 cells were infected with wild type parasites. Cells were treated with 500 µM IAA or the vehicle (EtOH) for 48 h. All wells were fixed and labeled with PanCp and Hoechst. The number of host cells (**a**) and *C. parvum* (**b**) in each well were imaged and counted on a Cytation 3 imager. **c** HCT-8 cells were infected with CpGP-mAID-TIR1 parasites. Cells were treated with 500 µM IAA or the vehicle (EtOH) for 24 h or 48 h. All wells were fixed and labeled with PanCp and Hoechst. The number of host cells (**c**) in each well was imaged and counted on a Cytation 3 imager. Each bar represents the mean ± SD for nine replicates in total from three experiments. Statistical analysis performed using was performed with a two-way ANOVA corrected for multiple comparisons by Sidak's method. All the comparisons between IAA and vehicle group showed no significant difference.
